# Translational suppression via IFG-1/eIF4G confers resistance to stress-induced RNA alternative splicing in *Caenorhabditis elegans*

**DOI:** 10.1101/2021.11.29.470369

**Authors:** Samantha C. Chomyshen, Cheng-Wei Wu

**Affiliations:** Department of Veterinary Biomedical Sciences, Western College of Veterinary Medicine, University of Saskatchewan, Saskatoon, SK, Canada; Toxicology Centre, University of Saskatchewan, Saskatoon, SK, Canada; Department of Biochemistry, Microbiology and Immunology, College of Medicine, University of Saskatchewan, Saskatoon, SK, Canada

## Abstract

Splicing of pre-mRNA is an essential process for dividing cells and splicing defects have been linked to aging and various chronic diseases. Environmental stress has recently been shown to alter splicing fidelity and molecular mechanisms that protect against splicing disruption remains unclear. Using an *in vivo* RNA splicing reporter, we performed a genome-wide RNAi screen in *Caenorhabditis elegans* and found that protein translation suppression via silencing of the conserved initiation factor 4G (IFG-1/eIF4G) protects against cadmium-induced splicing disruption. Transcriptome analysis of an *ifg-1* deficient mutant revealed an overall increase in splicing fidelity and resistance towards cadmium-induced alternative splicing compared to the wild-type. We found that the *ifg-1* mutant up-regulates >80 RNA splicing regulatory genes that are controlled by the TGF-β transcription factor SMA-2. The extended lifespan of the *ifg-1* mutant is partially reduced upon *sma-2* depletion and completely nullified when core spliceosome genes including *snr-1, snr-2*, and *uaf-2* are knocked down. Together, these data describe a molecular mechanism that provides resistance towards stress-induced alternative splicing and demonstrate an essential role for RNA homeostasis in promoting longevity in a translation-compromised mutant.

## INTRODUCTION

Post-transcriptional splicing of precursor mRNAs (pre-mRNA) is an essential step in the regulation of eukaryotic gene expression and contributes to proteome diversity (Baralle & Giudice 2017). Precision in RNA splicing is essential for maintaining cellular homeostasis, this includes alternative splicing of select transcripts into isoform variants in specific tissues, or in response to particular developmental cues to coordinate organismal growth (Nilsen & Graveley 2010; Wang et al. 2008). Errors in RNA splicing can disrupt cellular homeostasis, driven by the assembly of aberrant mRNA molecules that lead to the production of erroneous proteins with abnormal functions and detrimental consequences (Dutertre et al. 2011). A decline in RNA splicing fidelity has been reported during aging, and is prominently linked to many chronic human diseases including neurodegeneration and cancer (Tollervey et al. 2011; Scotti & Swanson 2015; Deschênes & Chabot 2017). However, the underlying cellular mechanism contributing to RNA splicing disruption remains poorly understood, and potential protective strategies have yet to be characterized. Uncovering mechanistic insights that protect splicing fidelity is intriguing as small molecules that modulate RNA splicing have been explored for their potential as chemotherapy agents (Eskens et al. 2013).

In the genetic model *Caenorhabditis elegans*, it was recently demonstrated that RNA splicing fidelity deteriorates with age and this dysregulation is hypothesized to be a central contributor to the decline in transcriptome and protein homeostasis (Heintz et al. 2017). Interventions that extend lifespan such as dietary restriction (DR) can improve splicing fidelity during aging; conversely, loss of splicing homeostasis nullifies DR-induced longevity, suggesting a co-dependency of DR and splicing homeostasis in extending lifespan. In Rhesus monkeys, caloric restriction extends lifespan and leads to modifications of RNA splicing as evident by the up-regulation of various spliceosome components at the transcriptome and proteome levels in the liver (Rhoads et al. 2018). This suggests that a causative relationship between RNA homeostasis and dietary restriction-induced longevity in *C. elegans* may be conserved in higher organisms. Next to DR, other interventions that significantly extend *C. elegans* lifespan include reduced insulin and TOR signaling (Kenyon et al. 1993; Robida-Stubbs et al. 2012), decreased mitochondrial respiration (Felkai et al. 1999; Feng et al. 2001), and inhibition of protein translation (Hansen et al. 2007; Pan et al. 2007); however, whether maintaining RNA homeostasis is essential for the longevity phenotype of these interventions remain to be characterized.

Recently, we showed that RNA splicing is disrupted in *C. elegans* upon exposure to the heavy metal cadmium (Wu et al. 2019). This demonstrated that environmental stress may promote aging by negatively influencing RNA homeostasis. In this study, we performed a genome-wide RNAi screen and found that inhibiting protein translation protects against cadmium-induced RNA splicing disruption. Focusing on the translation initiation factor *ifg-1* (initiation factor <u>4</u>G family), we show that reduced *ifg-1* improves RNA splicing fidelity and provides resistance against cadmium-induced alternative splicing. Suppression of *ifg-1* protects against cadmium-induced alternative splicing is dependent on the transcription factor *sma-2*, which regulates the expression of >80 genes involved in RNA splicing that is up-regulated in the *ifg-1* mutant. Depletion of *sma-2* and several core RNA splicing regulators reduced the lifespan of the *ifg-1* mutant, implicating an essential role for RNA splicing homeostasis in promoting longevity o translation compromised mutants. Overall, our results suggest a model where reduced protein translation up-regulates the expression of RNA splicing regulatory genes via the SMA transcription factors to improve RNA splicing fidelity under stress in promoting longevity.

## RESULTS

### Inhibiting protein translation protects against cadmium-induced alternative splicing

Exposure to environmental stress has been known to inhibit pre-mRNA splicing (Biamonti & Caceres 2009). We recently showed that the heavy metal cadmium can disrupt RNA splicing in *C. elegans,* potentially by altering snRNA processing function of the Integrator complex (Wu et al. 2019). To obtain a visual biomarker to study cadmium-induced RNA splicing disruption, we used an *in vivo* fluorescent alternative splicing reporter constructed with the *C. elegans ret-1* gene (Kuroyanagi et al. 2013). Briefly, this splicing reporter expresses a pair of fluorescent *ret-1* mini genes driven by the ubiquitous *eft-3* promoter. The mCherry reporter is modified with a +1 nucleotide insertion to exon 5, while the GFP reporter is modified with +1 insertion to exon 5 and +2 insertion to exon 6 (Fig. 1A). The nucleotide insertions alter the native reading frame of each fluorescent reporter, resulting in mCherry fluorescence when exon 5 is skipped or GFP fluorescence when exon 5 is included (Heintz et al. 2017; Kuroyanagi et al. 2013). Using this splicing reporter, we show that exposure to 300 µM of cadmium for 24 h results in a strong loss of GFP fluorescence, indicating an increased incidence of exon 5 skipping (Fig. 1A). To verify alternative splicing occurs to the endogenous *ret-1* gene, we exposed wild-type worms with 0, 100, 200, and 300 µM of cadmium followed by RNA extraction and PCR analysis with primer pairs that bind to exon 4 and exon 6 of the endogenous *ret-1* gene (Heintz et al. 2017). Two isoforms of the *ret-1* transcript are amplified and can be visualized via agarose gel electrophoresis, with the top band representing the full *ret-1* transcript expressing exons 4, 5, and 6, and the bottom band representing the skipped *ret-1* transcript expressing only exon 4 and 6 (Fig. 1B). Densitometry analysis showed that increasing cadmium exposure concentration leads to an increase in the relative skipped/full ratio of the *ret-1* gene (Fig. 1B). This verified that cadmium induces endogenous *ret-1* exon skipping.

**Figure 1.**
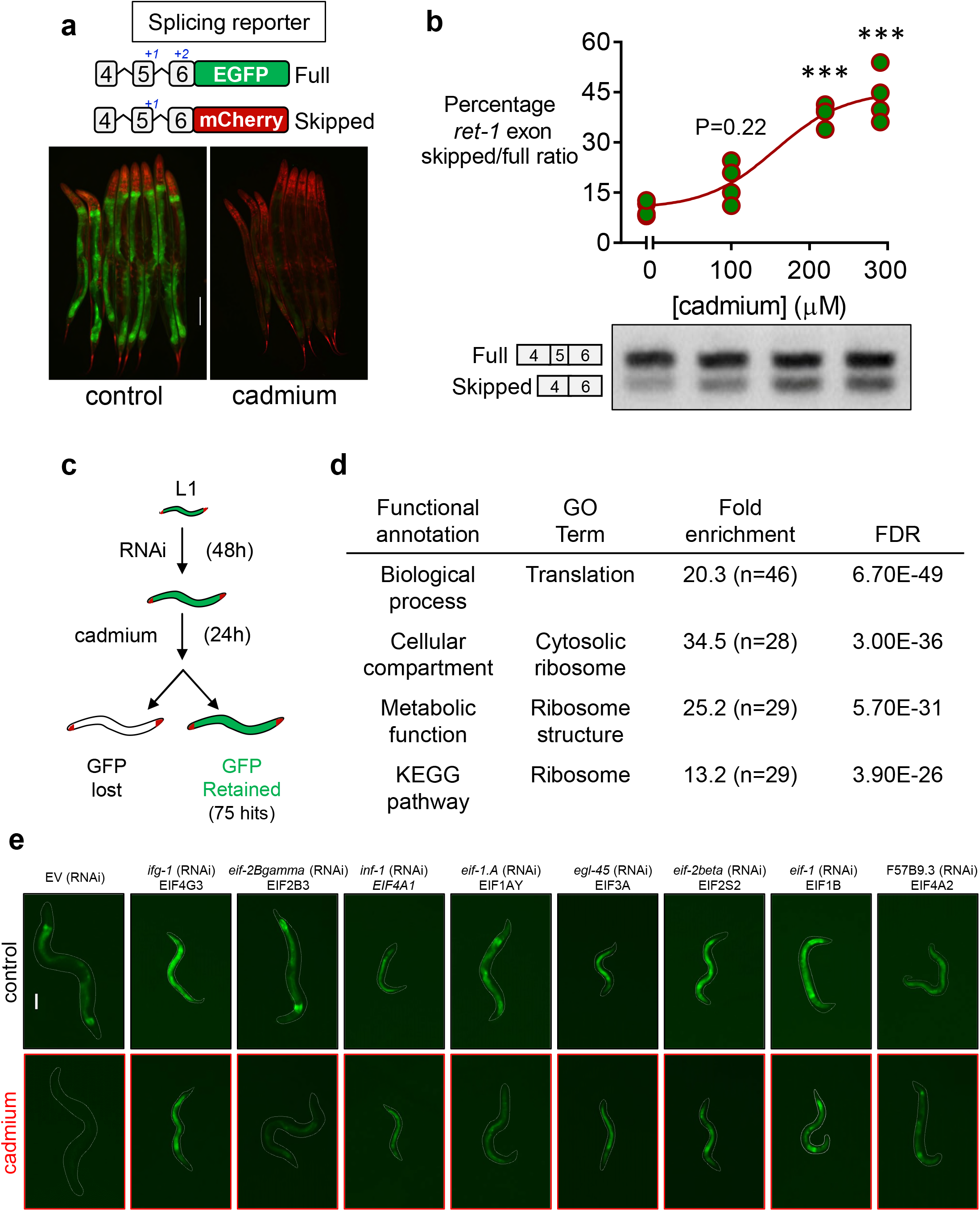
Suppression of protein translation protects against cadmium-induced exon skipping. **a.** Schematic of *in vivo ret-1* splicing reporter (Kuroyanagi et al. 2013; Heintz et al. 2017) and representative images of the splicing reporter under control condition and after cadmium exposure. Scale bar = 100 µm. **b.** Densitometry analysis and representative gel image of the relative ratio of skipped/full-length *ret-1* isoform after 0, 100, 200, and 300 µM of cadmium. N = 4 replicates with each RNA sample obtained from ∼200-300 worms. ***P<0.001 as determined via One-way ANOVA. **c.** Workflow of the genome-wide RNAi screen. Complete list of RNAi hits are shown in Table S1 **d.** DAVID enrichment analysis of genes encoding the 75 dsRNA clones that resulted in *ret-1* GFP retention after cadmium exposure. **e.** Representative images of *ret-1* GFP fluorescence in worms fed with dsRNA targeting translation initiation that strongly retained *ret-1* GFP fluorescence before and after cadmium exposure. Scale bar = 100µm.

We then performed a genome-wide RNAi screen to identify genetic regulators of cadmium-induced alternative splicing (Fig. 1C). We screened ∼19,000 dsRNA clones and identified 75 gene knockdowns that resulted in the retention of the *ret-1* GFP reporter fluorescence after cadmium exposure. Using DAVID functional analysis, we found a strong enrichment for protein translation processes among the 75 positive hits from the RNAi screen (Fig. 1D), with the majority of the RNAi targeting genes encoding small or large ribosomal proteins and various protein translation initiating factors (**Table S1**). Representative images demonstrating the effects of RNAi on the retention of *ret-1* GFP reporter fluorescence after cadmium exposure relative to EV control are shown in Fig. 1E. Overall, these findings revealed that suppression of protein translation protects against cadmium-induced alternative splicing.

### Suppression of *ifg-1* enhances *C. elegans* resistance to cadmium

It is well established that reduction of protein synthesis via depletion of various translation related genes extends lifespan in *C. elegans* (Pan et al. 2007; Hansen et al. 2007). We selected four genes from the RNAi screened that demonstrated high retention of *ret-1* GFP reporter fluorescence after cadmium exposure to determine their role in lifespan and cadmium resistance regulation. RNAi depletion initiated at day 1 of adulthood of *rps-23* (human homologue: ribosome protein S23), *ifg-1* (human homologue: eIF4G), F57B9.3 (human homologue: eIF4A2), and Y61A9LA.10 (human homologue: BMS1 ribosome biogenesis factor) all resulted in significant increases in mean lifespan of the worms compared to EV controls. Surprisingly, only RNAi depletion of *ifg-1* led to a consistent significant increase in cadmium survival relative to EV control (Fig. 2A; **Table S2**). As such, we focused our downstream analysis on *ifg-1* for further characterization.

**Figure 2.**
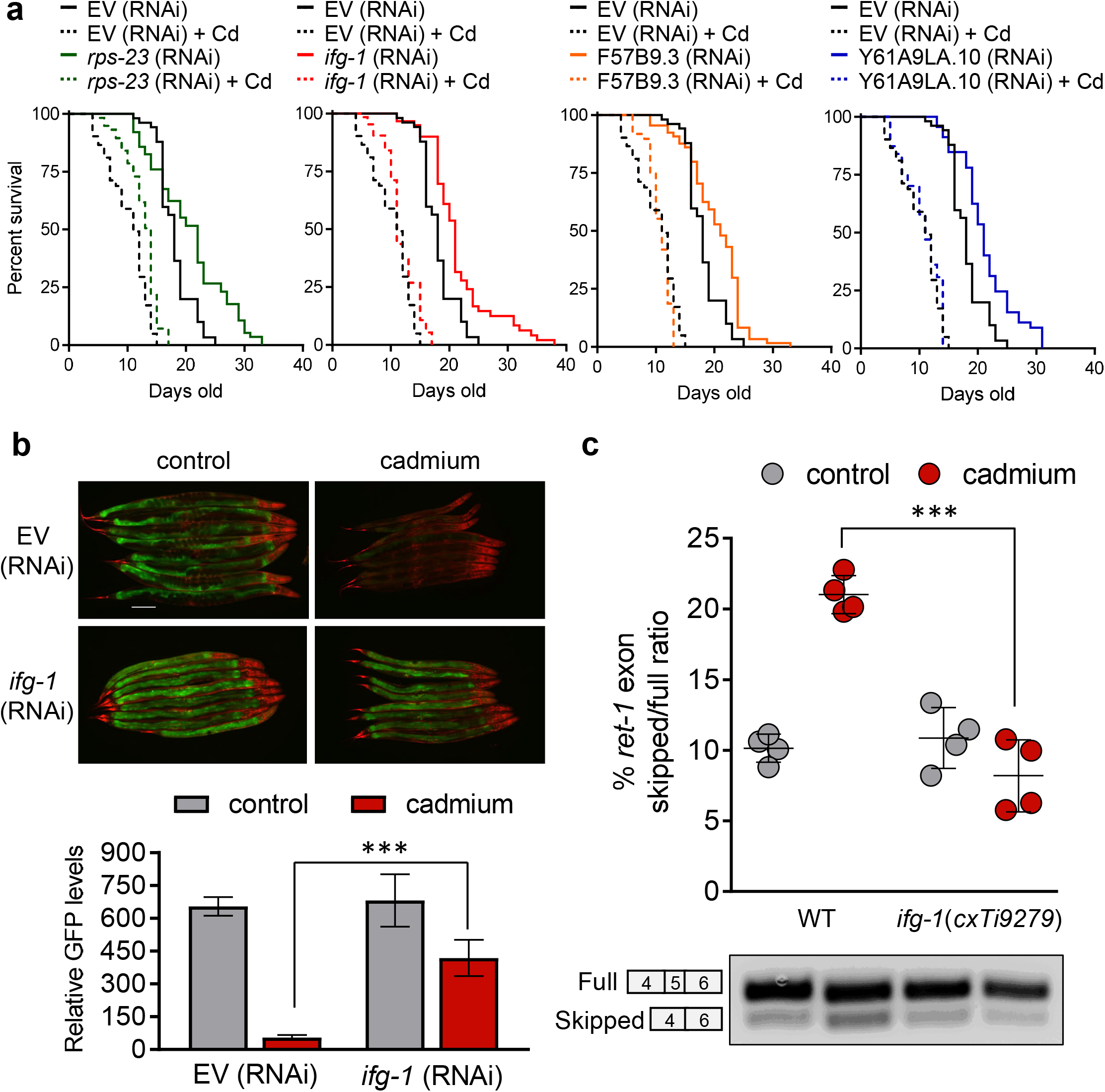
Reduced expression of *ifg-1* increases resistance towards cadmium toxicity. **a.** Lifespan and cadmium (Cd) survival of worms after *rps-23*, *ifg-1*, F57B9.3, and Y61A9LA.10 RNAi initiated at day 1 of adulthood. Three independent trials were performed for each assay, detailed statistics for individual trials are shown in Table S2. **b.** Representative image and quantification of *ret-1* GFP fluorescence between EV and *ifg-1* RNAi fed worms before and after cadmium exposure. N= 4 images each containing 8 worms for a total of 32 worms analyzed per condition. P<0.001 as determined via One-way ANOVA. **c**. Densitometry analysis and representative gel image of the relative ratio of skipped/full-length *ret-1* isoform in wild-type (WT) and *ifg-1(cxTi9279)* mutant worms before and after cadmium exposure. N = 4 replicates with each RNA sample obtained from ∼200-300 worms. Mean values are graphed with error bars representing standard deviation of the mean, ***P<0.001 as determined via One-way ANOVA.

Given that RNAi from the genome-wide screen was initiated at the L1 stage, knockdown of *ifg-1* resulted in the larval arrest of *C. elegans* at the L2/L3 stage. To verify that knockdown of *ifg-1* protects against cadmium-induced *ret-1* alternative splicing is not due to RNAi induced developmental delay, we first attempted to introduce a viable *ifg-1(cxTi9279)*partial loss of function mutation background into the *ret-1* splicing reporter strain (Morrison et al. 2014); however, we were not able to recover viable homozygous offspring (data not shown). Given that the *ret-1* splicing reporter has been shown to gradually lose a substantial amount of GFP fluorescence as early as 5 days old through aging (Heintz et al. 2017), it restricts the comparison of cadmium-induced reduction in GFP fluorescence to only young worms. To minimize developmental differences between EV and *ifg-1* RNAi within an age window where *ret-1* GFP is highly fluorescent, we initiated RNAi depletion of *ifg-1* at the L2/L3 stage for 48 hours followed by 24 hours of cadmium exposure. This ensured that comparisons were made when worms are 2 days old where *ret-1* GFP is still highly expressed. Worms with *ifg-1* depleted via RNAi at L2/L3 were able to reach adulthood and retained high levels of *ret-1* GFP fluorescence after cadmium exposure compared to the EV control (Fig. 2B). To verify that alternative splicing changes are also observed to the endogenous *ret-1* transcript, we extracted total RNA from wild-type and *ifg-1(cxTi9279)* worms with and without cadmium treatment. PCR analysis showed that cadmium exposure significantly increased the skipped/full ratio of the *ret-1* transcript in the wild-type, and this ratio remained unchanged in the *ifg-1(cxTi9279)* mutant (Fig. 2C). Overall, these findings revealed that reduction of protein translation via depletion of *ifg-1* increases *C. elegans* resistance to cadmium and protects against cadmium-induced alternative splicing.

### The transcriptome of *ifg-1* mutant mimics a cadmium stress response

To investigate how reduced *ifg-1* expression protects against cadmium toxicity, we first analyzed transcription changes in wild-type and *ifg-1(cxTi9279)* worms treated with and without cadmium by whole-transcriptome RNA sequencing (Fig. 3A). RNA-sequencing revealed that *ifg-1(cxTi9279)* worms show a 57% reduction in the *ifg-1* mRNA level (Fig. 3B), this is comparable to a previous study indicating a 42% decrease in *ifg-1* mRNA in this mutant as determined via Northern blot (Morrison et al. 2014). The *ifg-1(cxTi9279)* worms displayed a dramatically altered transcriptome compared to the wild-type, with 1,423 up-regulated genes and 4,246 down-regulated genes (>2-fold change, Padj<0.05; **Table S3**. KEGG pathway analysis revealed that *ifg-1(cxTi9279)* up-regulated genes enriched to antioxidant stress response pathways and metabolism, while down-regulated genes enriched to pathways regulating neuroactivity along with calcium and Wnt signaling pathways (Fig. 3C). Up-regulation of stress response genes in *ifg-1* depleted worms was similarly demonstrated previously through translation state array analysis showing an increased abundance of stress genes associated with highly translated ribosome fractions (Rogers et al. 2011).

**Figure 3.**
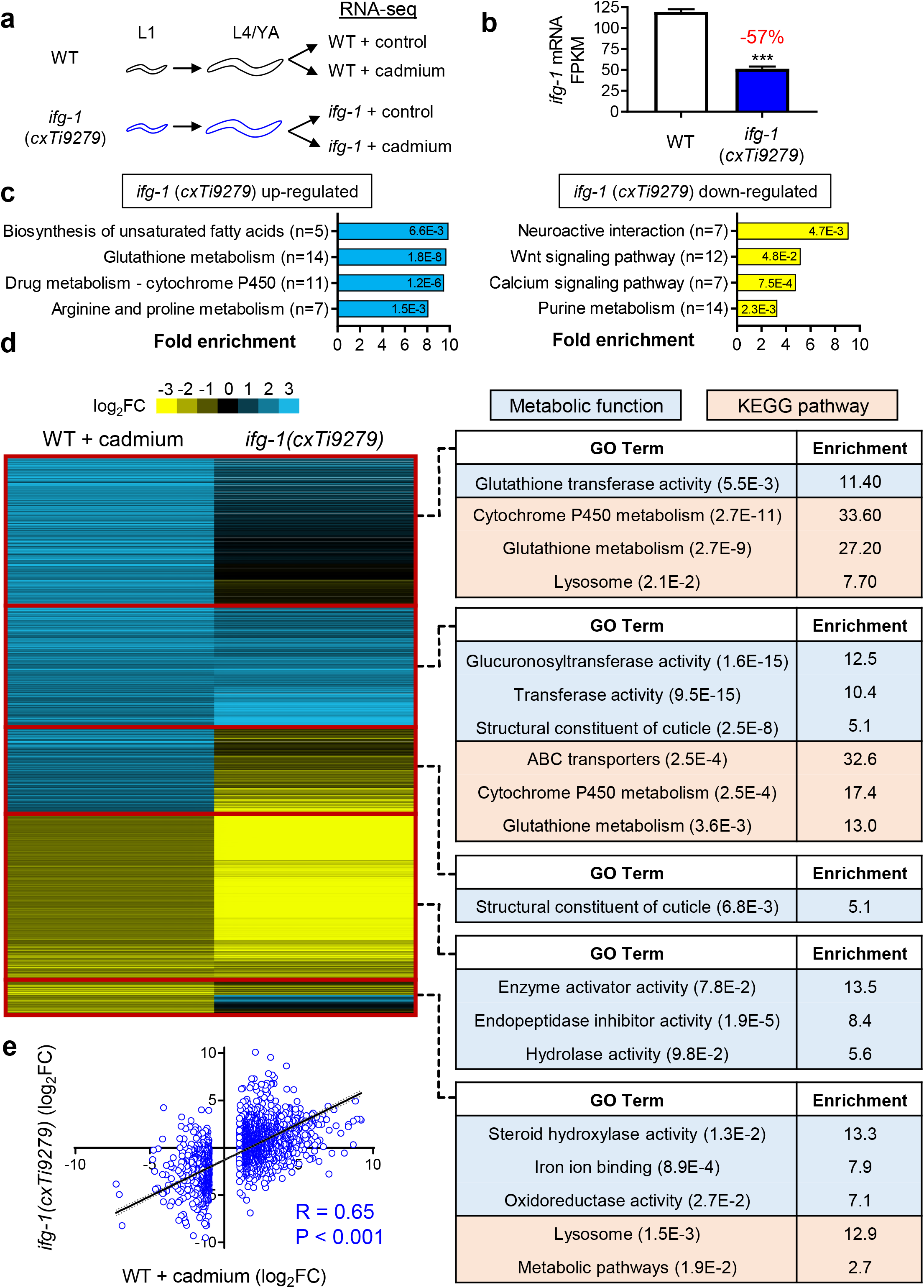
RNA-sequencing reveals transcriptome similarities between wild-type cadmium exposure and *ifg-1* mutant. **a.** Experimental workflow for RNA extraction for transcriptome sequencing. **b.** FKPM value of *ifg-1* mRNA in wild-type (WT) and *ifg-1(cxTi9279)* mutant. Mean values are graphed with error bars representing standard deviation of the mean; N = 3 replicates with each RNA sample obtained from ∼2,000–3,000 worms, ***P<0.001 as determined via student t-test. **c.** KEGG pathway enrichment of gene up or down-regulated by >2-fold (FDR<0.05) in the *ifg-1(cxTi9279)* mutant. **d.** Heat map of differentially expressed genes in WT + cadmium and *ifg-1(cxTi9279)* mutant worms cluster by relative expression change, all genes were differentially expressed by >2-fold (FDR<0.05). DAVID enrichment analysis of genes within each heat map segment determined based on relative similarity and difference in expression between WT + cadmium and *ifg-1(cxTi9279)* mutant. Enriched metabolic function and KEGG pathways are shown. **e.** Linear regression analysis of genes with >2-fold change (FDR<0.05) in WT + cadmium compared to the expression of the corresponding gene in *ifg-1(cxTi9279)* mutant. R indicates Pearson correlation value with P<0.001 as determined by the linear regression *F* test.

A clustered heat map analysis demonstrated that gene expression changes to the *ifg-1(cxTi9279)* mutant were highly similar to wild-type after cadmium exposure (Fig. 3D). Genes that were up-regulated in both *ifg-1(cxTi9279)* mutant and wildtype after cadmium exposure enrich specifically to metabolic functions and KEGG pathways involved in antioxidant detoxification including cytochrome P450 metabolism, glutathione activity, and ABC transporters. Meanwhile, genes down-regulated in both conditions enrich to metabolic enzyme functions including endopeptidase inhibitor and hydrolase (Fig. 3D). For cadmium-induced gene expression changes greater than 2-fold, linear regression analysis revealed a highly significant (P<0.001) Pearson correlation coefficient of +0.65 when compared to expression changes to the same genes in the *ifg-1(cxTi9279)* mutant (Fig. 3E). These results demonstrated that cadmium and *ifg-1(cxTi9279)* mutant share overlapping changes to the transcriptome, highlighted by activation of the antioxidant stress response.

### Cadmium-induced alternative splicing is modified in the *ifg-1* mutant

Through RNA-sequencing, we next compared the transcriptome-wide alternative splicing of the *ifg-1(cxTi9279)* mutant relative to wild-type worms treated with cadmium. This is accomplished by first determining the inclusion level (IncLevel) of a given transcript (Fig. 4A) followed by their relative differences between experimental conditions (IncLevel Difference) to determine significant alternative splicing events (Shen et al. 2014). Cadmium exposure in wild-type worms resulted in 437 significant alternative splicing events (183 IncLevel Difference > 0, 254 IncLevel Difference <0) across 5 different categories of alternative splicing (Fig. 4B). Analysis of the same 437 alternative splicing events revealed that only 93/183 (IncLevel Difference > 0) and 135/254 (IncLevel Difference < 0) alternative splicing events are statistically significant in the *ifg-1(cxTi9279)* mutant (Fig. 4B). Next, we performed a linear regression analysis of IncLevel difference between cadmium treated *ifg-1(cxTi9279)* and cadmium treated wild-type for the 437 alternative splicing events and found a negative Pearson correlation of −0.37 (P<0.001) (Fig. 4C). This suggested that the relative degree of cadmium-induced alternative splicing changes observed in wild-type worms is regressed in the *ifg-1(cxTi9279)* mutant. Correlation analysis for each alternative splicing type revealed that while four out of five categories showed a negative correlation coefficient, only exon skipping and alternative 3’ splice site (A3SS) coefficients are statistically significant (Fig. S1). It should be noted that exon skipping and A3SS account for 73% of all alternative splicing events induced by cadmium (320/437), and the lack of statistical significance may be due to a relatively small number of splicing events in other categories. Next, we analyzed the relative percentage of intron and intergenic reads between the wild-type and *ifg-1(cxTi9279)* mutant. Given that the cDNA sequencing library was constructed with poly-A capture enrichment, these two parameters are indicators of aberrant splicing and have previously been shown to increase with age and after spliceosome inhibition (Heintz et al. 2017). In the *ifg-1(cxTi9279)* mutant, the relative percentage of both intron reads and intergenic reads are significantly lower compared to the wild-type, suggesting an increase in RNA splicing fidelity (Fig. 4D).

**Figure 4.**
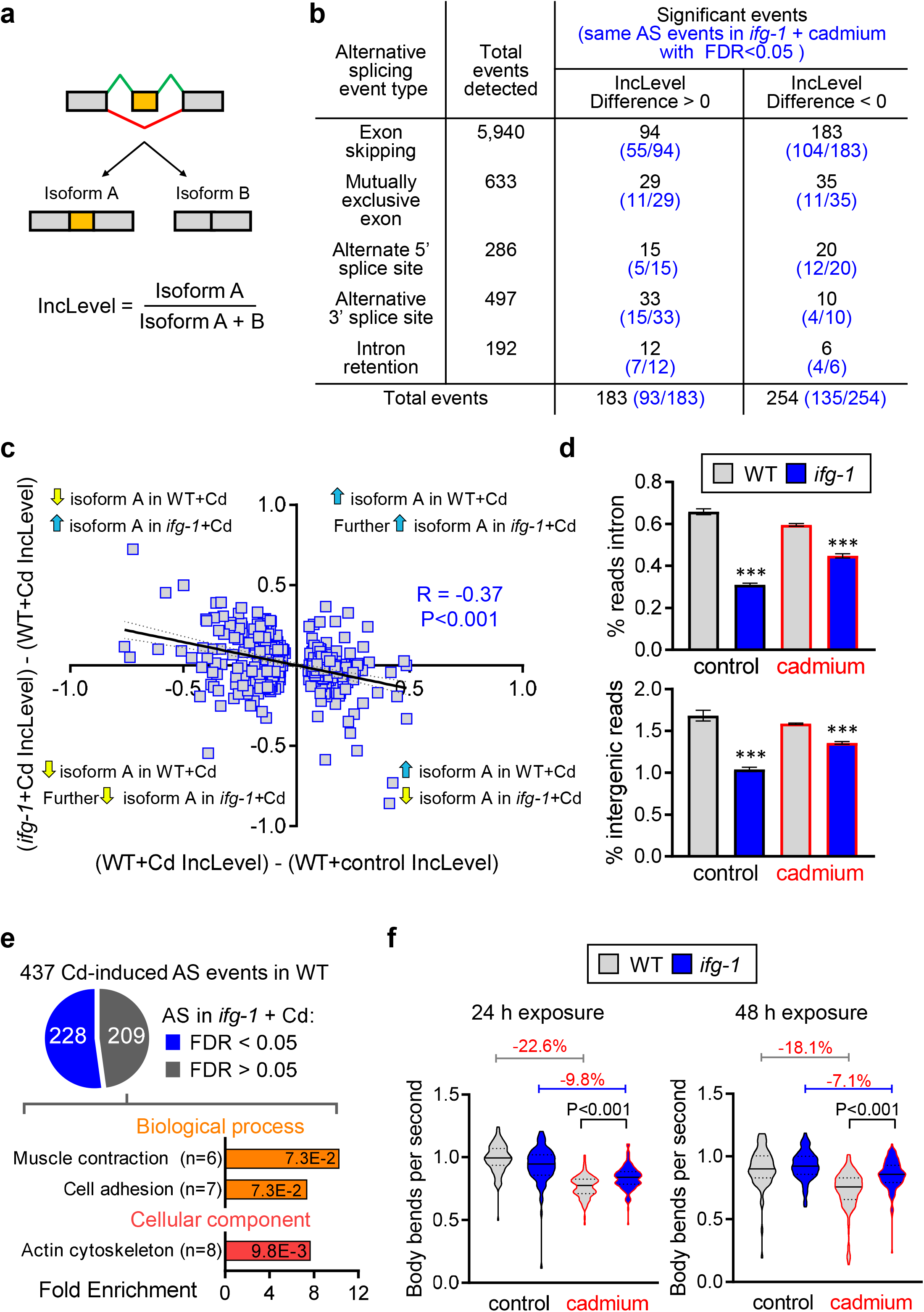
*ifg-1* mutant protects against cadmium-induced alternative splicing. **a.** Schematic of exon skipping event produces two transcript isoforms used to calculate inclusion level by rMATS, adapted from (Shen et al. 2014). **b.** The number of alternative splicing events detected in wild-type (WT) worms after cadmium (Cd) exposure is categorized by different event types. Same alternative splicing events were evaluated for statistical significance in *ifg-1(cxTi9279)* worms after cadmium exposure and presented in the bracket. **c.** Linear regression analysis of inclusion level difference between *ifg-1(cxTi9279)* and WT for the 437 significantly altered alternative splicing events detected in cadmium exposed wild-type worms. **d.** Percentage of intron and intergenic region reads in WT and *ifg-1(cxTi9279)*before and after cadmium exposure. Mean values are graphed with error bars representing standard deviation of the mean; N = 3 replicates with each RNA sample obtained from ∼2,000–3,000 worms, ***P<0.001 as determined by One-way ANOVA compared to WT. **e.** DAVID enrichment analysis of 209 genes that are not alternatively spliced in *ifg-1(cxTi9279)* after cadmium exposure. Enriched biological processes and cellular compartments are shown. **f.** Swim motility of WT and *ifg-1(cxTi9279)* worms after 24 h or 48 h cadmium treatment. Mean values are graphed with error bars representing the standard deviation of the mean. N = 89-130 worms scored for 24 h and 77-140 worms for 48 h in each condition, combined from two independent trials. P<0.001 as determined by One-way ANOVA.

Functional analysis of the 209 genes accounting for 48% of cadmium-induced alternative splicing observed in the wild-type that are not significantly altered in the *ifg-1(cxTi9279)* mutant revealed enrichment towards biological functions related to muscle contraction and cell adhesion located within the actin cytoskeleton cellular compartments (Fig. 4E). Given that cadmium-induced alternative splicing of genes functioning in muscle regulation, we measured the swimming behavior of wild-type and *ifg-1(cxTi9279)* mutant after 24 h and 48 h of cadmium exposure. Cadmium exposure for 24 h led to a 22.6% decrease in wild-type swimming motility as determined by body bends per second, this is significantly lower compared to a 9.8% decrease observed in the *ifg-1(cxTi9279)* mutant (Fig. 4F). Similar trends were also observed after 48 h cadmium exposure. Together, these results revealed that cadmium-induced alternative splicing observed in wild-type worms are reduced in the *ifg-1(cxTi9279)* mutant and that these alterations may provide protection against cadmium-induced impairment to muscle functions.

### SMA transcription factors regulate *ifg-1* mediated gene expressions

To gain mechanistic insights on how reduced *ifg-1* provides resistance against cadmium-induced alternative splicing, we performed a small-scale RNAi screen to identify transcription factors that are required to maintain *ret-1* GFP fluorescence after *ifg-1* depletion and cadmium exposure (Fig. 5A). We screen ∼500 dsRNA clones targeting *C. elegans* transcription factors and identified 2 dsRNA clones corresponding to *sma-2* and *sma-3* that suppressed *ifg-1* induced retention of *ret-1* GFP following cadmium exposure (Fig. 5B). The *C. elegans sma-2* and *sma-3* encode receptor-regulated Smad proteins within the TGF-β/BMP signaling pathway that was initially characterized for its role in body size regulation (Savage et al. 1996; Savage-Dunn & Padgett 2017). Given the defined role of SMA proteins functioning as transcription factors, we compared the transcriptome of a *sma-2(rax5)* mutant previously described with a defect in lipid metabolism to the transcriptome of the *ifg-1(cxTi9279)* mutant from this study (Yu et al. 2017). We observed a striking overlap between genes that are up-regulated in the *ifg-1(cxTi9279)*mutant but down-regulated in the *sma-2(rax5)* mutant (FDR>0.05, no FC cut off) (Fig. 5C), with 79% (2,899/3,680) of genes down-regulated in the *sma-2(rax5)* mutant found to be up-regulated in the *ifg-1(cxTi9279)* mutant. Functional analysis of the 2,899 overlapped genes revealed enrichment towards RNA metabolism pathways including RNA transport and the spliceosome (Fig. 5D). Other KEGG pathways enriched include proteasome, nucleotide excision repair (NER), and ER processing. Analysis of the two RNA-sequencing datasets revealed moderate up-regulation (1.2 to 1.9-fold) of 88 genes in the *ifg-1(cxTi9279)* mutant functioning in various processes in RNA splicing regulation, which were down-regulated in the *sma-2(rax5)* mutant (Fig. 5E). We then performed linear regression analysis on the expression of 5,840 genes that are significantly up or down-regulated (FDR<0.05) in both *ifg-1(cxTi9279)* and *sma-2(rax5)*mutants and observed a Pearson correlation coefficient of −0.41 (P<0.001) (Fig. 5F). While no RNA-sequencing data was available for a *sma-3* mutant, similar result was also observed when the transcriptome of a *sma-4(rax3)* mutant was compared to *ifg-1(cxTi9279)*, showing an overlap of 3,699 genes and a Pearson correlation coefficient of −0.52 (P<0.001) (Fig. S2). Overall, our results demonstrated that knockdown of *sma-2* suppressed *ifg-1(cxTi9279)* mediated resistance towards cadmium-induced alternative splicing, and this may be attributed to the requirement for *sma-2* in regulating transcription of RNA splicing regulatory genes that are up-regulated in the *ifg-1(cxTi9279)* mutant.

**Figure 5.**
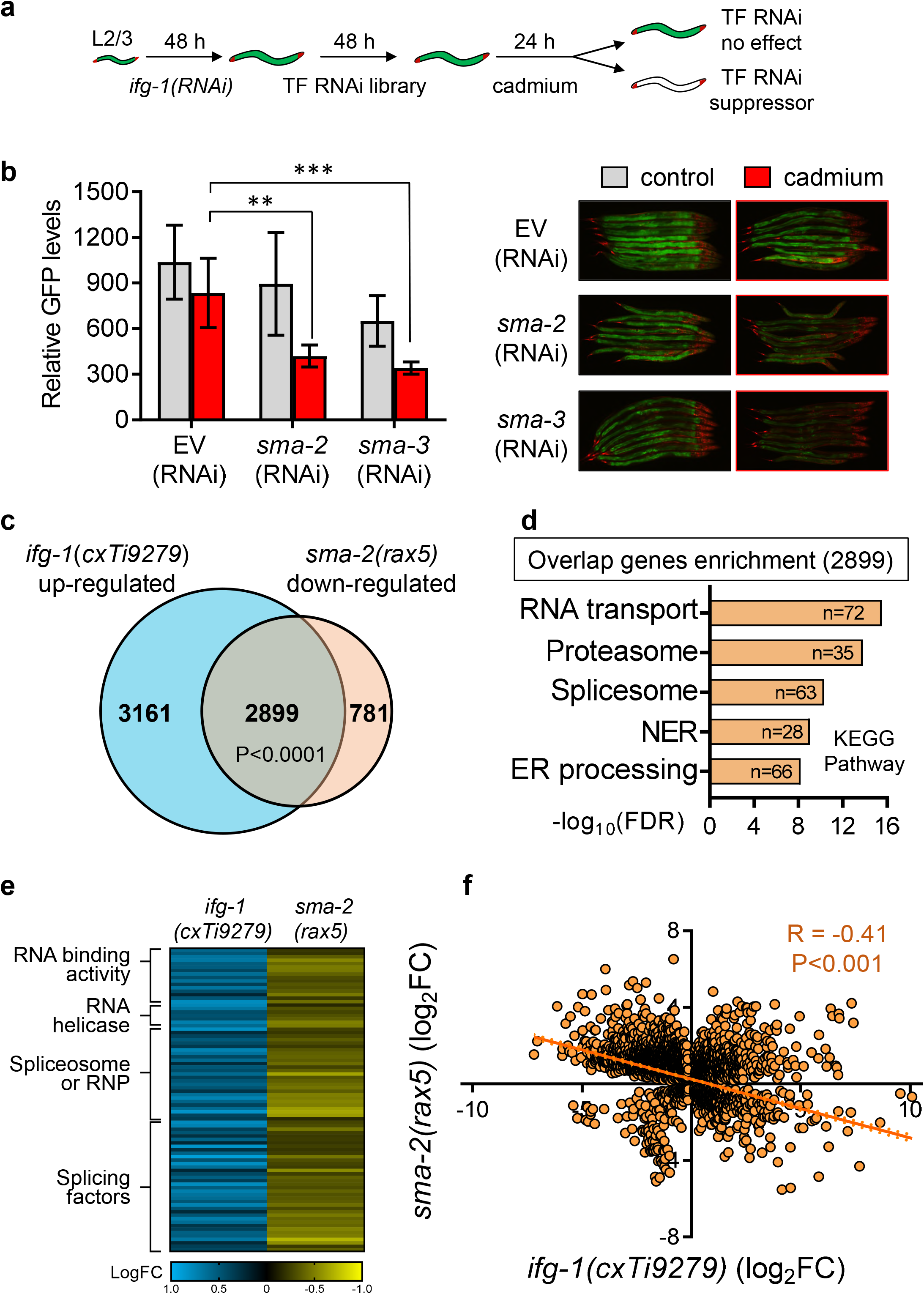
*sma* regulates transcription genes altered in the *ifg-1* mutant. **a.** Workflow of the transcription factor sub-library RNAi screen. **b.** Quantification and representative image and of *ret-1* GFP fluorescence between EV, *sma-2*, and *sma-3* RNAi-fed worms before and after cadmium exposure. Mean values are graphed with error bars representing the standard deviation of the mean. N= 4 images each containing 8 worms for a total of 32 worms analyzed per condition. **P<0.01 and ***P<0.001 as determined via One-way ANOVA. **c.** Venn diagram illustrating number of genes overlapping between *ifg-1(cxTi9279)* up-regulated and *sma-2(rax5)* down-regulated. P<0.001 as determined by Fisher’s exact test. Raw transcriptome data for *sma-2(rax5)* were obtained from (Yu et al. 2017). **d.**KEGG pathway enrichment of overlapped genes between *ifg-1(cxTi9279)* up-regulated and *sma-2(rax5)* down-regulated. Heat map of 88 RNA splicing regulatory genes significantly up-regulated in *ifg-1(cxTi9279)* and down-regulated in *sma-2(rax5)*. Linear regression analysis of fold change for genes differentially expressed in *ifg-1(cxTi9279)* compared to its corresponding fold change in *sma-2(rax5)*. R indicates Pearson correlation value with P<0.001 as determined by the linear regression *F* test.

### Core RNA splicing regulatory genes are required for *ifg-1* longevity

Our data suggest that *sma* and RNA splicing factors function downstream of reduced *ifg-1*. To test this, we used RNAi to knock down *sma-2, sma-3*, and four RNA splicing regulatory genes *snr-1*, *snr-2*, *uaf-2*, and *rsp-2*to determine their effects on *ifg-1(cxTi9279)* lifespan. The mean lifespan of *ifg-1(cxTi9279)* is significantly increased by ∼25% compared to the wild-type, this is consistent with previous reports (Fig. 6A-B; **Table S4, S5**) (Rogers et al. 2011; Pan et al. 2007). Knockdown of *sma-2* and *sma-3* slightly reduced *ifg-1(cxTi9279)* lifespan by 7% and 8% respectively (Fig. 6A-B; P<0.05 in 2/3 trials for *sma-2* and 3/3 trials for *sma-3*; **Table S4**), but did not reduce the lifespan of wild-type worms. We next investigated whether four RNA splicing regulatory genes up-regulated in the *ifg-1(cxTi9279)* mutant are required for its longevity phenotype. RNAi knockdown of small nuclear ribonucleoproteins *snr-1* and *snr-2* beginning at day 1 of adulthood significantly reduced wild-type lifespan by an average of 15.8% and 18.5% respectively, and completely nullified the long-lived phenotype of the *ifg-1(cxTi9279)* mutant with *snr-1*RNAi causing a 27.2% decrease (P<0.001) and *snr-2*RNAi causing a 33.2% decrease (P<0.001) in mean lifespan (Fig. 6C, D; **Table S5**). Interestingly, knockdown of the U2 small nuclear RNA auxiliary factor *uaf-2* beginning at day 1 of adulthood also completely nullified *ifg-1(cxTi9279)* mutant’s long-lived lifespan (22% decrease in mean lifespan, P<0.001) but did not affect wild-type lifespan (Fig. 6E; **Table S5**). Meanwhile, RNAi knockdown of the serine arginine-rich protein *rsp-2* did not affect either the wild-type or *ifg-1(cxTi9279)* mutant lifespan (Fig. 6F; **Table S4**). Together these experiments demonstrated that *sma-2* and *sma-3* are partially required for *ifg-1(cxTi9279)* mutant’s long-lived lifespan whereas core RNA splicing regulatory factors including *snr-1*, *snr-2*, and *uaf-2* are completely essential for longevity.

**Figure 6.**
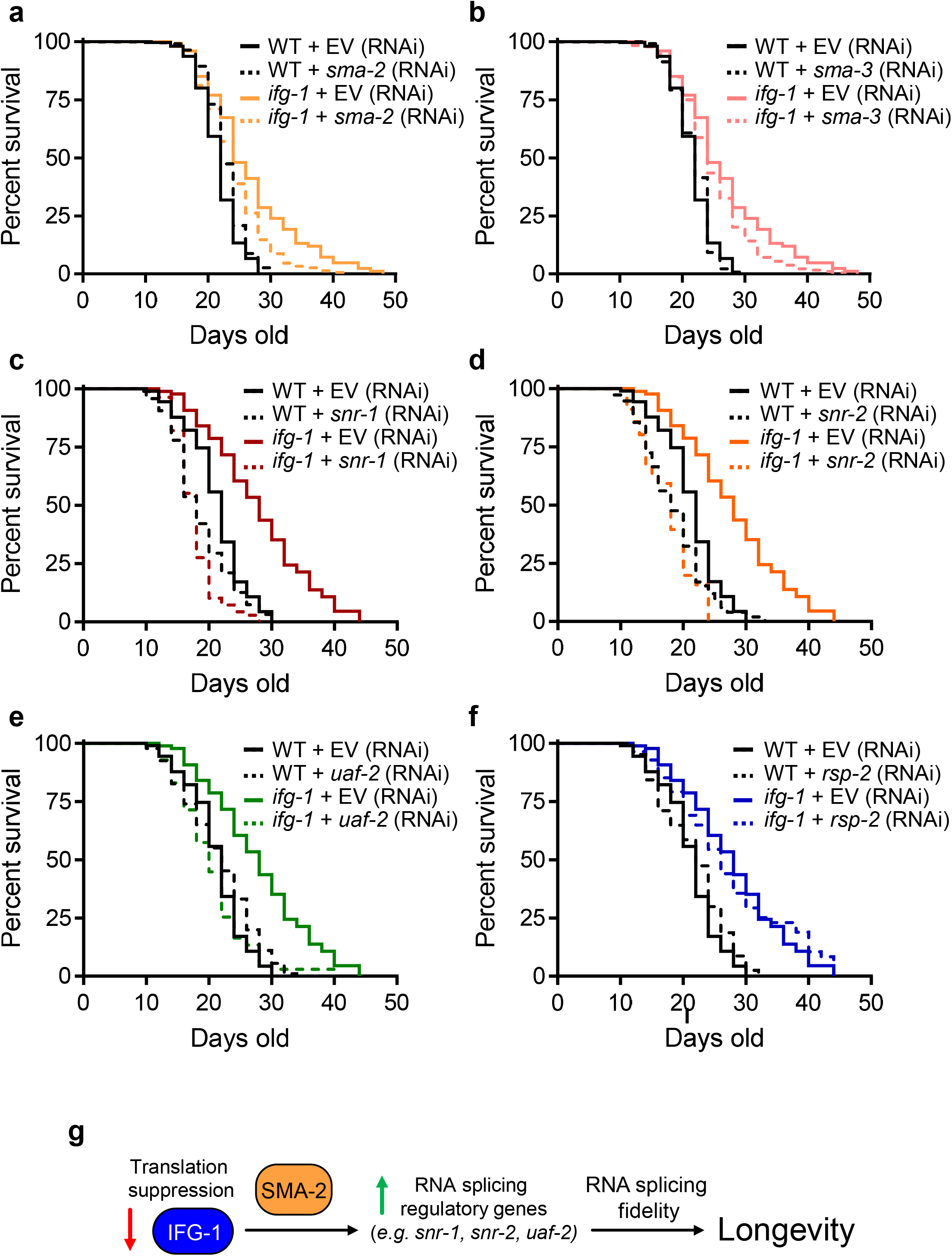
Essential RNA splicing regulator genes are required for the longevity of the *ifg-1* mutant. Lifespan wild-type (WT) and *ifg-1(cxTi9279)* mutant worms after **a.** *sma-2* (RNAi), **b.** *sma-3* (RNAi), **c.** *snr-1* (RNAi), **d.** *snr-2* (RNAi), **e.** *uaf-2* (RNAi), and **f.** *rsp-2* (RNAi). Three independent trials were performed for each assay, detailed statistics for individual trials are shown in Table S4 and S5.

## DISCUSSION

RNA splicing is an essential step in the regulation of eukaryotic gene expression and is a major contributor to transcriptome and proteome diversity (Dutertre et al. 2011). Aberrant RNA splicing is an underlying factor in many human diseases including cancer and neurodegeneration (Tollervey et al. 2011; Scotti & Swanson 2015; Deschênes & Chabot 2017). Recently, it was demonstrated that RNA splicing fidelity declines with age in *C. elegans* suggesting a causative link between RNA splicing homeostasis and healthy aging (Heintz et al. 2017). To date, only handful of studies have examined the effects of environmental stress on RNA splicing homeostasis (Yost & Lindquist 1986; Biamonti & Caceres 2009; Dutertre et al. 2011; Wu et al. 2019), and molecular mechanisms that protect against stress-induced RNA splicing disruption have not been described yet. Here, we show that protein translation suppression in *C. elegans* via reduced *ifg-1* protects against cadmium-induced alternative splicing (Fig. 1-4), and we propose that this mechanism signals through the SMA family of transcription factors to regulate the expression of RNA splicing regulatory genes in maintaining splicing fidelity to promote longevity (Fig 5-6, Fig. 6g).

### Influence of translational suppression on RNA splicing and aging

It is well established that reduced protein translation via mutation or knockdown of various ribosomal and translation factors can significantly extend lifespan in worm, fly, and yeast (Curran & Ruvkun 2007; Hansen et al. 2007; Pan et al. 2007; Rogers et al. 2011; Kapahi et al. 2004; Zid et al. 2009; Kaeberlein et al. 2005; Steffen et al. 2008). Mechanistically, a conserved cellular response to reduced protein translation includes the selective up-regulation and increase in translation of various genes involved in stress signaling (Rogers et al. 2011; Zid et al. 2009). Consistently, suppression of protein translation in worms provides enhanced resistance to various environmental stress including hypoxia, thermotolerance, oxidative stress, heavy metal stress, and ER stress (Anderson et al. 2009; Howard et al. 2016; Wang et al. 2010). The mechanism behind lifespan extension induced by protein translation suppression is likely to be multifactorial, given that transcriptome-wide alteration to gene expression is observed (Fig. 3), and presumably, to the proteome as well. It is hypothesized that the combination of enhanced stress resistance and global reduction to protein translation both contribute to the longevity phenotype (Steffen & Dillin 2016). Enhanced stress resistance may provide an increase in cellular repair response to mitigate the progressive buildup of molecular damage accumulated through aging (Gems & Partridge 2013). Our results show that the *ifg-1* mutant up-regulates various stress responsive genes involved in xenobiotic detoxification (Fig. 3). This is consistent with a previous report showing enhanced oxidative stress resistance in *ifg-1* depleted worms, which is dependent on the SKN-1 transcription factor that functions as a master regulator of antioxidant defense (Wang et al. 2010; An et al. 2005; Wu et al. 2017). Interestingly, while loss of *skn-1* abolishes *ifg-1*’s oxidative stress resistance, it had minimal impact on its longevity (Wang et al. 2010), suggesting that enhanced oxidative stress response may not be the primary factor in promoting longevity in translation compromised mutants.

Meanwhile, reduced translation may directly decelerate the proposed hypertrophic effect of protein synthesis in later life, this is supported by evidence that translation fidelity decreases with age leading to an increase in accumulation of translational errors (Anisimova et al. 2018; Gems & Partridge 2013). Mild inhibition of translation has also been shown to increase protein folding accuracy in mammalian cells, suggesting a positive effect of reduced translation on enhancing overall protein synthesis fidelity (Meriin et al. 2012). Similar to translation, RNA splicing fidelity also decline with age, a deterioration that can be protected through dietary restriction (DR) in *C. elegans* (Heintz et al. 2017). In this study, we provide evidence that translation inhibition via reduced *ifg-1* also protects RNA splicing fidelity (Fig. 5), though this may be accomplished via independent mechanisms compared to DR that did not show global changes to splicing factor mRNA expression (Fig. 5E). Expression of splicing regulators has been shown to be a determinant factor in mouse and human longevity (Lee et al. 2016), and down-regulation of splicing factors is observed with increasing chronological age in man (Holly et al. 2013). In calorically restricted Rhesus monkeys that lived on average 20% longer than the control group, transcriptome analysis of liver biopsy samples showed up-regulation of various components of the spliceosome (Rhoads et al. 2018). Though it should be cautioned that the correlative relationship between splicing factor expression and aging likely differs between organisms and can vary across different tissues, and functional studies are required to determine their relative contribution to aging. In this study we showed that depletion of select core RNA splicing factors (*i.e. snr-1*, *snr-2*, and, *uaf-2*) nullify lifespan extension observed in *ifg-1* mutants, suggesting that maintaining RNA splicing homeostasis is essential for longevity facilitated by reduced translation.

Why does translational suppression lead to enhanced RNA splicing fidelity? Under periods of environmental stress, cells naturally cope by suppressing global protein translation via eIF2α phosphorylation (Wek et al. 2006). This allows cells to redirect energy resources to prioritize other essential needs, which may include enhancing RNA processing to reduce potential stress-induced disruption to splicing fidelity. While cells can protect against mis-spliced mRNA molecules via RNA surveillance mechanisms such as nonsense-mediated decay (NMD), costs associated with degradation of an erroneous transcript is energetically expensive given that it cost at least 10 ATP molecules to perform splicing of each intron, and that NMD relies on the activity of various ATP dependent hydrolysis reactions (Lynch & Marinov 2015; Serdar et al. 2016). Increasing RNA splicing fidelity also reduces the likelihood that an erroneous transcript is translated into an altered protein, which can cost on average 100-200 molecules of ATP to degrade (Lynch & Marinov 2015; Peth et al. 2013). Given that the combined energetic cost of RNA transcription and splicing are magnitudes lower than the cost of protein translation, increasing splicing fidelity may serve as a protective mechanism during periods of stress to ensure the production of accurate mature transcripts prior to translation to avoid loss of energy investment towards the synthesis and degradation of aberrant proteins.

### SMA signaling in aging

TGF-β/Sma signaling is a critical developmental pathway in *C. elegans* initially discovered for its role in regulating body size growth and male tail morphogenesis (Savage-Dunn & Padgett 2017; Savage et al. 1996). In the last decade, a role for TGF-β/Sma signaling in aging has also been described. Loss of function mutations to *sma* genes delay reproductive aging characterized by an extended offspring production window supported by improved oocyte and germline quality (Luo et al. 2009; Luo et al. 2010). In the oocyte of *sma-2* mutants, reproductive aging is delayed via transcriptional up-regulation of genes involved in cell cycle regulation, DNA damage repair, and cell death programming, while down-regulating genes involved in oviposition, proteolysis and cell differentiation (Luo et al. 2010). Recently, a role for SMA in regulating lipid metabolism has also been described, with *sma-2* and *sma-4* mutants transcriptionally up-regulating various β-oxidation genes to reduce lipid storage in *C. elegans* (Yu et al. 2017; Clark et al. 2018). These transcriptional programs appear to be distinct from the regulation of genes involved in RNA metabolism described in this study, suggesting a pleiotropic role for the TGF-β/Sma signaling in regulating diverse cellular functions. In our study, we show that depletion of either *sma-2* or *sma-3* does not shorten wild-type lifespan, but partially reduced the longevity of the *ifg-1* mutant. This suggests a differential requirement of *sma* genes in influencing lifespan that is dependent on the genetic background, where SMA may function as critical transcription factors in long-lived mutants to regulate the expression of genes required to potentiate longevity. This genotype-specific requirement for *sma* genes in longevity was also recently demonstrated in an insulin signaling mutant where loss *sma-3* partially reduced the extended lifespan of the *daf-2(e1370)*worms (Clark et al. 2021).

In summary, we have described a new mechanism through which inhibition of protein translation protects against stress-induced alternative splicing in *C. elegans*. We show that reducing the translation initiation factor *ifg-1* up-regulates the expression of RNA splicing regulators, increases RNA splicing fidelity, and protects against stress-induced alternative splicing. Overall, these findings reveal novel mechanistic insights into RNA splicing regulation under stress and highlight the importance of maintaining RNA homeostasis in promoting longevity.

## EXPERIMENTAL PROCEDURES

### C. elegans strains

*C. elegans* strains were grown and maintained at 20°C using standard methods (Brenner 1974). The following strains were used: wild-type N2 Bristol, KH2235 *lin-15(n765)* ybIs2167[*eft-3::ret-1E4E5(+1)E6-GGS6-mCherry+eft-3::ret-1E4E5(+1)E6(+2) GGS6-EGFP+lin-15(+)+pRG5271Neo*]X, *ifg-1(cxTi9279)*.

### Genome-wide RNAi screen and RNAi experiments

RNAi was performed as described previously by feeding *C. elegans* strains of engineered *E. coli [HT115(DE3)]* that transcribed double-stranded RNA (dsRNA) homologous to a target gene (Wu et al. 2016; Wu et al. 2019). Approximately 19,000 dsRNA from the ORFeome RNAi feeding library (Open Biosystems, Huntsville, AL) and the MRC genomic RNAi feeding library (Geneservice, Cambridge, UK) were used for screening. Briefly, synchronized L1 larvae of KH2235 *C. elegans* obtained from the alkaline hypochlorite method were grown in liquid nematode growth medium (NGM) with dsRNA producing bacteria for 2 days, followed subsequently by exposure to 300 µM cadmium chloride for 24 h and manually screened for relative fluorescence of the *ret-1* GFP reporter with an Olympus SZX61 stereomicroscope. Clones that resulted in the retention of *ret-1* GFP signal after 24 h cadmium exposure were noted and rescreened three additional times for verification. All subsequent RNAi experiments were performed on NGM agar plates in the presence of 50 µg mL^-1^ carbenicillin and 100 µg mL^-1^ isopropyl β-D-thiogalactopyranoside (IPTG). *E. coli* expressing the pPD129.36 (LH4440) plasmid encoding 202 bases of dsRNA that are not homologous to any predicted *C. elegans* gene, or referred to as empty vector (EV), was used as an RNAi control.

### RNA extraction and PCR

Total RNA was extracted with the Invitrogen Purelink RNA mini kit (ThermoFisher, 12183020) according to the manufacturer’s instructions using a QSonica Q55 sonicator. RNA was extracted from four biological replicates per condition, with each sample containing 200-300 synchronized worms. RNA samples were treated with DNAse I (ThermoFisher, EN0521) before cDNA synthesis using the Invitrogen Multiscribe^TM^ reverse transcriptase (ThermoFisher, 4311235). Analysis of *ret-1* alternative splicing was carried out using primers previously described with the Applied Biosystems ProFlex thermocycler (Heintz et al. 2017). PCR product was separated on a 2.5% agarose gel and stained with SYBR Green I nucleic acid gel stain (ThermoFisher, S7563) for band visualization using a BioRad Gel Doc EQ system; ImageJ (NIH) was used to quantify the relative band intensity.

### Fluorescent microscopy

Preparation of *C. elegans* for fluorescent imaging was carried out as previously described (Murray et al. 2020). Fluorescent images were captured with a Retiga R3 camera mounted to either a Zeiss Axioskop 50 or an Olympus SZX61 stereomicroscope. GFP fluorescence was detected using filter sets with Ex: 460-495 nm / Ex: 510-550 nm and RFP fluorescence was detected using filter sets with Ex: 540-580 nm / Em: 610 nm. Grayscale images were converted to color images using ImageJ (Schneider et al. 2012). Worm images that were used to quantify relative fluorescence were prepared by mounting eight worms per slide immobilized by 2% sodium azide on a 2% agarose pad. Relative fluorescence was calculated in ImageJ using the Measure function. Background signal was subtracted from each image by defining an area on the same image where fluorescence signal was absent, with the dimension of the defined area constant for all images. Composite images displaying both GFP and RFP fluorescence were created with the Merge Channel function in ImageJ.

### Lifespan and cadmium survival assay

Lifespan assays were performed at 20°C on NGM RNAi plates seeded with HT115 *E. coli* with the EV dsRNA plasmid used for control. To circumvent the larval arrest phenotype of most RNAi tested in this study, synchronized L1 worms were first grown on EV dsRNA until day 1 of adulthood, after which the worms were transferred to RNAi plates seeded with the corresponding dsRNA to initiate knockdown of the gene of interest. For two dsRNA that did not affect larval development (*sma-2, sma-3*), RNAi was initiated at the L1 stage. Lifespan assays were performed without the use of FUDR with progeny separation accomplished via daily picking during the worm’s reproductive window. For cadmium survival assay, synchronized L1 worms were first grown on EV dsRNA until day 1 of adulthood followed by transfer to RNAi plates seeded with the corresponding dsRNA for 48 h to initiate gene knockdown. Day 3 old worms were then transferred to corresponding RNAi seeded NGM plates containing 300 µM of cadmium chloride. For both lifespan and survival assays, worms were considered dead if they did not respond to gentle prodding with a metal pick and censored when they displayed protruding gonad or vulva. Survival assays were scored daily while lifespan assays were scored every 2 days. Three independent trials were performed for each assay with the number of animals used in each experiment described in the corresponding supplementary table.

### Whole-transcriptome sequencing and data analysis

Wild-type and *ifg-1(cxTi9279)* worms were synchronized at the L1 stage and grown on EV dsRNA NGM plates until adulthood followed by transfer to EV dsRNA NGM plates with or without 300 µM of cadmium chloride for 24 h. Total RNA from wild-type and *ifg-1(cxTi9279)* worms with and without cadmium treatment were extracted from three biological replicates for each condition with ∼2,000-3,000 worms per replicate. RNA samples were sent to Novogene (Sacramento, CA) on dry ice for cDNA library preparation with oligo(dT) enrichment followed by RNA-sequencing and statistical analysis. HISAT2 (v2.0.5) was used to map sequencing reads to the WBcel235 genome, with FeatureCounts (v1.5.0-p3) used for gene quantification, DESeq2 (v1.20.0) for differentially analysis, and rMATS (v3.2.5) for alternative splicing analysis (Kim et al. 2019; Anders & Huber 2010; Shen et al. 2014). Gene ontology enrichment was performed using DAVID functional analysis (Huang et al. 2009). Heat maps showing differentially expressed genes were analyzed using Gene Cluster 3.0 with uncentered average linkage and visualized with Java TreeView (v1.2.0). Raw sequence data and processed data are deposited to the NCBI Gene Expression Omnibus with the following accession number GSE184491. Raw data from previously published RNA-sequencing of *sma-2(rax5)* and *sma-4(rax3)*were retrieved from the NCBI Sequence Read Archive with the BioProject ID PRJNA395096 (Yu et al. 2017). The Galaxy (usegalaxy.org) open-source platform was used to analyze differential gene expression data using the methods described above (Afgan et al. 2018).

### Measurement of motility

Wild-type and *ifg-1(cxTi9279)* worms were synchronized at the L1 stage and grown on EV dsRNA NGM plates until adulthood followed by transfer to EV dsRNA NGM plates with or without 300 µM of cadmium chloride for 24 h or 48 h. *C. elegans* swimming rate (body bends per second in liquid) was measured by re-suspending worms grown on a 6-cm NGM agar plate with ∼5 mL of M9 buffer. Worms were allowed to acclimate to the liquid condition for at least 1 minute, after which the swimming behavior was recorded by a Retiga R3 camera mounted to an Olympus SZX61 stereomicroscope for 30 seconds. The video file was processed with the wrMTrck program in ImageJ to calculate body bends per second (Nussbaum-Krammer et al. 2015).

### Statistical analyses

Graphical data and statistical analysis were generated using the GraphPad Prism software (v7.04). For statistical analysis, student’s t-test was performed when two means were compared, one-way ANOVA with Tukey’s correction was used for multiple comparisons with one factor, and two-way ANOVA with Sidak correction used for multiple comparisons with two factors. F-test for linear regression was used to determine statistical significance when comparing relationships between two sets of RNA-sequencing data, Fisher’s exact test was used to determine statistical significance between Venn diagram overlap genes. For lifespan and survival analysis, the OASIS2 software (https://sbi.postech.ac.kr/oasis2/) was used to calculate mean lifespan, 95% confidence interval, and statistical significance between conditions using the Log-rank test (Kim et al. 2016). For all statistical tests *P<0.05, **P<0.01, and ***P<0.001, FDR correction was used for all P values involving RNA-sequencing datasets.

## ACKNOWLEDGEMENT

Some *C. elegans* strains were provided by the *Caenorhabditis* Genetic Centre (University of Minnesota, Minneapolis, MN) which is supported by the NIH Office of Research Infrastructure Programs (P40 OD010440). We thank Dr. Hidehito Kuroyanagi (Tokyo Medical and Dental University) for providing the KH2235 strain and Dr. Brett Keiper (East Carolina University) for providing the *ifg-1(cxTi9279)* strain. This work was supported by an NSERC Discovery Grant (04486) and an SHRF Establishment Grant (4971) to CWW. SCC was supported by a WCVM graduate teaching fellowship award.

## AUTHOR CONTRIBUTIONS

CWW designed the study, performed experiments, and analyzed data. SCC performed the majority of the experiments and analyzed the results. CWW and SCC wrote the manuscript.

## DATA AVAILABILITY

All datasets supporting this manuscript are found within the article and its supplementary files. RNA-sequencing data generated from this study (raw and annotated) are available on the NCBI GEO data repository GSE184491.

## SUPPORTING INFORMATION

**Figure S1.**
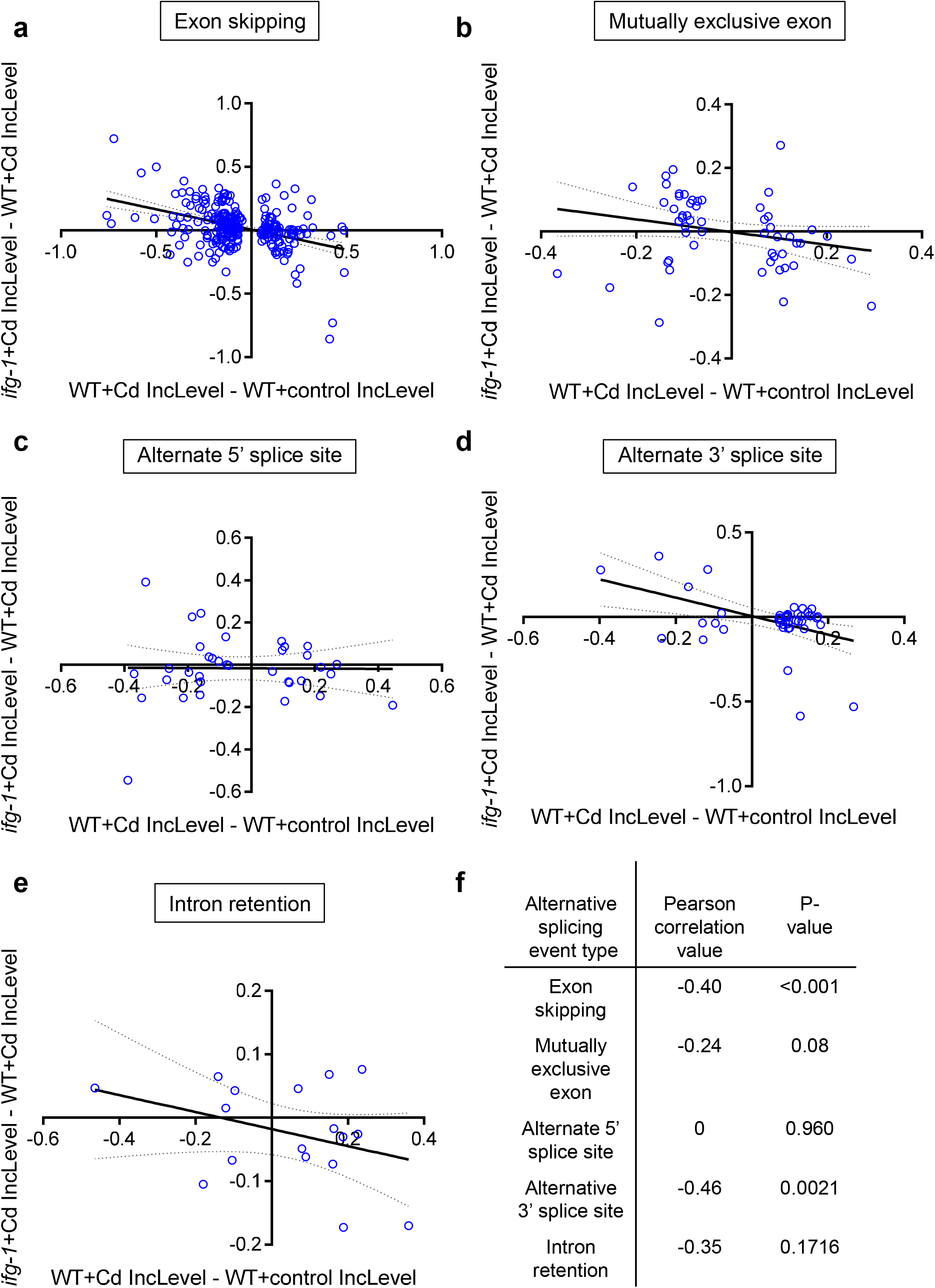
Linear regression analysis of five alternative splicing types. IncLevel difference between *ifg-1(cxTi9279)* and wild-type worms for the 437 cadmium-induced alternative splicing events categorized based on **a.** exon skipping, **b.**mutually exclusive exon, **c.** alternative 5’ splice site, **d.**alternative 3’ splice site, and **e.** intron retention. **f.** Correlation coefficient and P-value analysis of inclusion level difference between *ifg-1(cxTi9279)* and wild-type categorized by different alternative splicing events.

**Figure S2.**
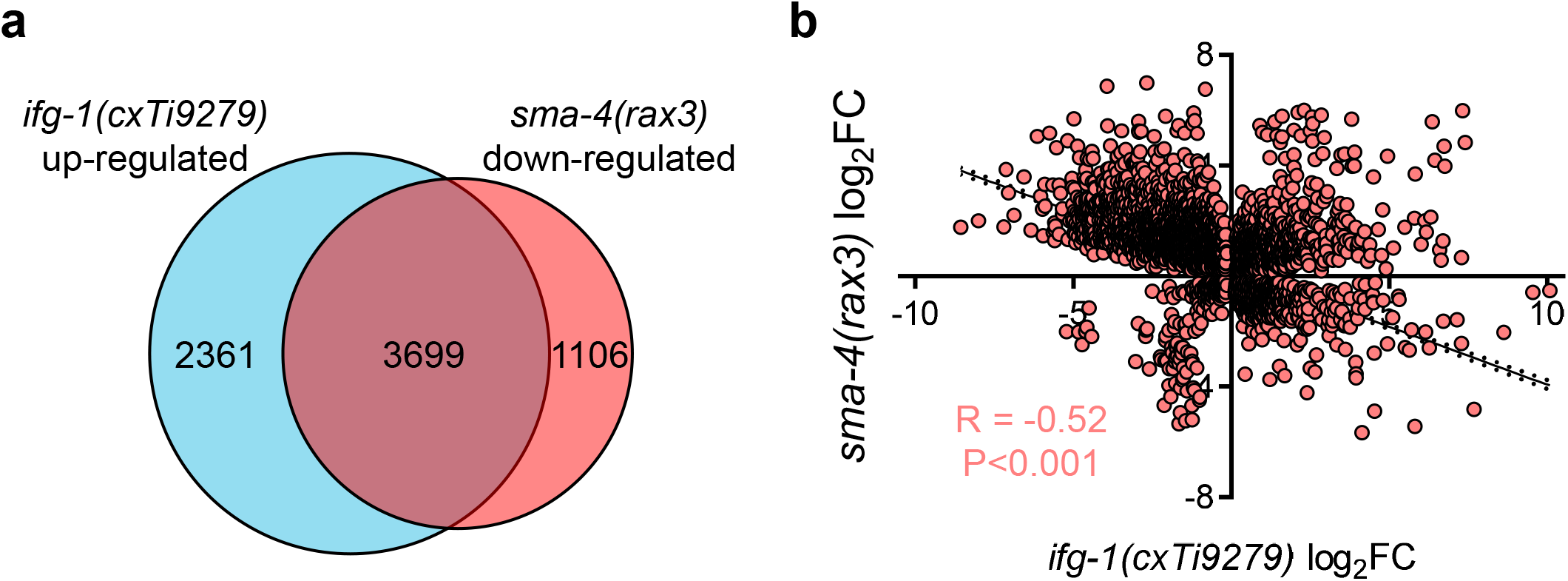
*sma-4* regulates *ifg-1(cxTi9279)* up-regulated genes. **a.** Venn diagram illustrating number of genes overlapping between *ifg-1(cxTi9279)* up-regulated and *sma-4(rax3)* down-regulated. P<0.001 as determined by Fisher’s exact test. Raw transcriptome data for *sma-4(rax3)* were obtained from (Yu et al. 2017) **b.** Linear regression analysis of fold change for genes differentially regulated in *ifg-1(cxTi9279)* compared to its corresponding fold change in *sma-4(rax3)*. R indicates Pearson correlation value with P<0.001 as determined by the linear regression *F* test.

**Table S1.** Genome-wide RNAi hit list

**Table S2.** Longevity and cadmium survival statistic for *rps-23, ifg-1*, F57B9.3, and Y61A9LA.10 RNAi in wild-type trials.

**Table S3.** Annotated RNA-sequencing results

**Table S4.** Longevity statistic for *sma-2* and *sma-3* RNAi in wild-type and *ifg-1(cxTi9279)* mutant trials.

**Table S5.** Longevity statistic for *snr-1*, *snr-2*, *uaf-2*, and *rsp-2* RNAi in wild-type and *ifg-1(cxTi9279)* mutant trials.

